# Polytraumatic SCI worsens maladaptive plasticity in spinal motor systems

**DOI:** 10.64898/2026.06.25.734362

**Authors:** Jason H Gumbel, Jacob A Davis, Kerui Gong, Cleopa Omondi, Jeffrey Sacramento, Emma G Iorio, Abel Torres-Espin, Jenny Haefeli, Kazuhito Morioka, Adam R Ferguson, J. Russell Huie

## Abstract

Spinal cord injury (SCI) results in dysfunction of both motor and sensory systems, which can be characterized by neuropathic pain, hypersensitivity, muscular spasticity and rigidity. Most SCIs result from incidents such as vehicle accidents or falls, resulting in polytraumatic SCI that includes peripheral injuries in addition to direct CNS damage. Recent findings suggest that spinal cord synaptic plasticity plays a crucial role in neuropathic pain pathophysiology, specifically in association with spinal sensitization and the consequent onset of AMPA-related maladaptive plasticity. Further findings have demonstrated that nociceptive peripheral stimulation in the acute phase of SCI results in maladaptive spinal synaptic plasticity by overdriving GluA2-lacking calcium-permeable AMPARs (CP-AMPARs). Here, we investigated the effect of a spared nerve injury (SNI) in conjunction with SCI to determine the effect of polytraumatic SCI on maladaptive plasticity in the spinal cord. Near-IR quantitative Western blot analysis demonstrated that SCI+SNI increases spinal GluA1 expression, but not GluA2. Patch-clamp confirmed that AMPAR currents in spinal motorneurons increase after SCI with SNI, and decrease after the administration of NASPM, a CP-AMPAR antagonist. Data-driven analysis using non-linear principal components analysis (NL-PCA) also demonstrated that SCI with SNI produces a multivariate signature of AMPAR plasticity that is observed in other forms of nociceptive peripheral input, indicating a general mechanism for maladaptive plasticity in spinal motor systems in response to polytraumatic SCI.

## Introduction

Spinal cord injury (SCI) is a condition characterized by profound impairment, affecting around 276,000 survivors in the United States alone. This results in an estimated $40 billion in annual financial impact from health care costs and lost productivity.^1,2^ SCI is often accompanied by a high incidence of concomitant polytraumatic peripheral injuries, including bone fractures, lacerations, and abrasions.^3–5^ Initially, these injuries may be viewed as insignificant compared to the spinal injury itself. However, the nociceptive input to the spinal cord that these injuries produce has lasting detrimental effects that inhibit the functional recovery of sensory and motor systems (maladaptive plasticity).^6,7^ Preclinical work on rodent SCI models has demonstrated that a peripheral nociceptive stimulus results in lasting alterations in neurons located in the dorsal horn of the spinal cord through mechanisms similar to hippocampal learning/memory.^7,8^ Indeed, concomitant peripheral injuries below the site of SCI as well as the consequent additional immobility, are ubiquitous in a clinical SCI setting,^9–13^ with polytraumatic SCI associated with lower American Spinal Injury Association (ASIA) Impairment Scale (AIS) at the time of hospital discharge, and additionally significantly increases hospital stay after initial injury.^14^

Outside the context of SCI, disrupted signal transduction and peripheral injury alter synaptic strength in the central nervous system (CNS), resulting in plasticity that inhibits functional recovery and is characterized by increased incidence of chronic pain and hyperreflexia or spasticity, called maladaptive plasticity.^7,15–21^ There is evidence that SCI causes an increase in postsynaptic localization of GluA2-lacking, calcium permeable AMPA receptors (CP-AMPARs) on motorneurons.^22,23^ AMPARs are well known to act a part within dynamic changes in learning and memory within the healthy CNS.^24–27^ More recently, AMPARs have been implicated within the pathological CNS, contributing to the negative behavioral effects described above.^28–35^ Over-activation of AMPA receptors is implicated in a host of CNS disorders, including traumatic brain injury^36^, epilepsy^37,38^, and stroke.^39,40^ Within experimental CNS injury, it has been shown that surgical incisions and bone fractures can drive increases in glutamate toxicity.^15,41^

The role of these cellular phenomena must be further explored within the area of spinal cord disorders. What is known is that changes in AMPAR subunits on lower motorneurons are associated with excitotoxicity and cell dysfunction in the hyperacute phase of SCI and amyotrophic lateral sclerosis.^22,42,43^ Previously, our laboratory has placed particular emphasis on AMPAR subunit structure in determining the form of plasticity within the injured spinal cord, implicating them in both maladaptive plasticity that undermines recovery after injury and adaptive plasticity that promotes good neural health.^20,22,28,44^ The structure of these AMPAR subunits at the synapse is determined in part by peripheral input. Widely, research in this area has modeled this input using peripheral nociceptive stimulation, stressful stimuli, and concomitant injuries intended to mimic polytraumatic afferent input.^45–49^ In particular, electrical noxious stimulation of the tail in SCI rats, forced hindlimb unloading, and peripheral injury models have demonstrated a shift towards hyper-excitable, GluA2-lacking, and calcium permeable AMPA receptors.^8,22,23,42^

Here we use a spared nerve injury, a common model of peripheral injury,^50–52^ to test the role of synaptic spinal AMPAR activity after polytraumatic SCI. Using quantitative protein expression analyses, spinal motorneuron electrophysiology, and multivariate machine learning, we present findings suggesting that polytraumatic SCI results in CP-AMPAR mediated maladaptive changes within spinal motor circuitry.

## Results

### Peripheral spared nerve injury after SCI drives GluA1 but not GluA2 expression in spinal synaptoneurosomes

To assess the effect of peripheral injury on spinal AMPA receptor expression, we paired a spinal transection injury with a peripheral nerve injury model in which the sural nerve is spared (**Figure 1A**). We then applied a quantitative near-infrared western blot to synaptoneurosomal-enriched lumbar spinal tissue (**Figure 1B**). Antibodies for GluA1 and GluA2 were first measured on a 1:2 total protein dilution curve to determine the optimal linear range for detection, ensuring parametric western blot results (**Figure 2A**).^53^ This procedure yielded a wide linear range for both GluA1 and GluA2. Assessment of AMPAR protein in ventral lumbar synaptoneurosomal spinal tissue demonstrated a significant increase in GluA1 in response to SCI+SNI, compared to SCI alone (**Figure 2B**). A concomitant increase in GluA2 was not observed. These findings indicate that the addition of SNI to a spinal injury increases AMPAR expression at synaptic sites, and that the differential expression of GluA1 and GluA2 suggests that this difference reflects an increase in GluA2-lacking, calcium-permeable AMPA receptors after peripheral injury.

**Figure 1.**
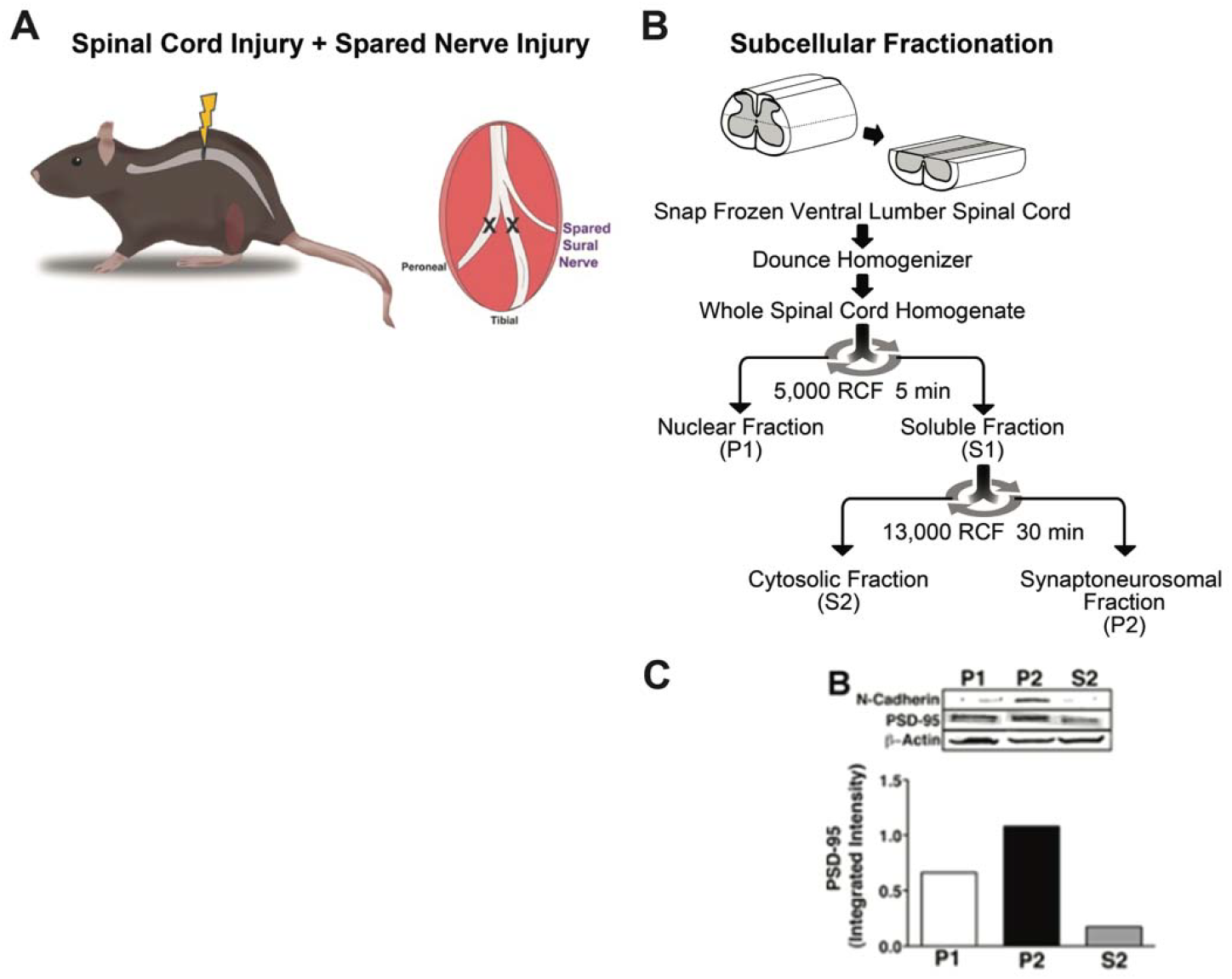
Experimental model and spinal cord preparation for biochemical assay. A) Model of spared nerve injury (SNI). First, a complete spinal transection was performed at the T9 spinal level. Immediately after closing the surgical site, the sciatic nerve was exposed using blunt dissection of the leg muscles. Both the peroneal and tibial nerves were ligated and cut, while the sural nerve was undisturbed. The surgical site was then closed, and tissue collection took place 15 minutes after the SNI surgery. B) Biochemical methods to rigorously and reproducibly assess synaptic plasticity in the ventral horn after peripheral stimulation. Cords were snap frozen in liquid nitrogen within 5 min of decapitation, and stored at −80°C. (A) On the day of the assay, the cord was warmed to −20°c and the lumbar enlargement (~1cm) was processed to isolate plasma membrane and synaptoneurosomal compartments. (C) The P2 fraction is characterized by enrichment in the plasma membrane protein N-cadherin and modest synaptic (PSD-95) enrichment.

**Figure 2.**
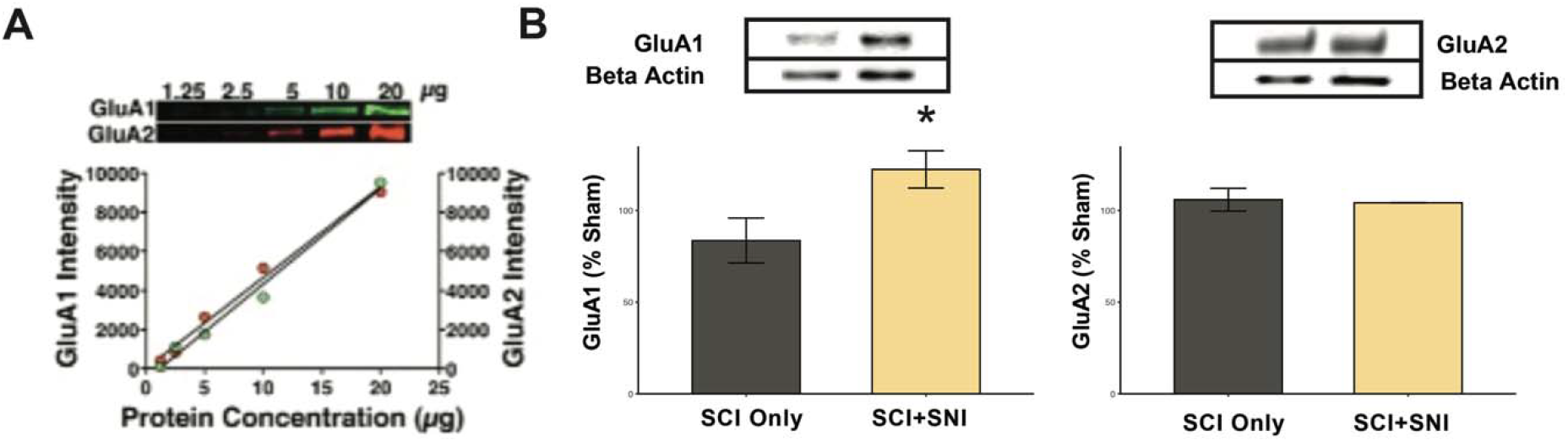
Spared nerve injury after SCI induces differential expression of spinal synaptic AMPAR subunits. (A) Serial protein dilutions were used to determine the linear range for near-infrared western blot analysis of GluA1 and GluA2 subunits, ensuring a parametric association between western band fluorescent intensity and target protein concentration. (B) Assessment of GluA1 and GluA2 AMPAR subunits in ventral spinal tissue of the lumbar enlargement (expressed as percent relative to the double sham group) showed a significant GluA1 increase in the SCI+SNI group compared to the SCI only group (* p < 0.05). No concomitant increase in GluA2 subunit expression was observed (p > 0.05). Mean ± SEM.

### Peripheral spared nerve injury after SCI increases CP-AMPAR currents in spinal neurons and increases the response to AMPA

To further investigate the electrophysiological function of ventral horn spinal neurons, we conducted several patch-clamp experiments. We started by recording AMPA-induced currents in motorneurons from a sham group. Most neurons were located in layers VIII – IX based on Rexed laminae. Fast puff application of AMPA (100 uM, 1 s) induced a quick inward current (**Figure 3A**). The specificity of currents was confirmed by the almost complete elimination of currents by cyanquixaline (CNQX) bath application, which is an AMPAR antagonist (**Figure 3B-C**).

**Figure 3.**
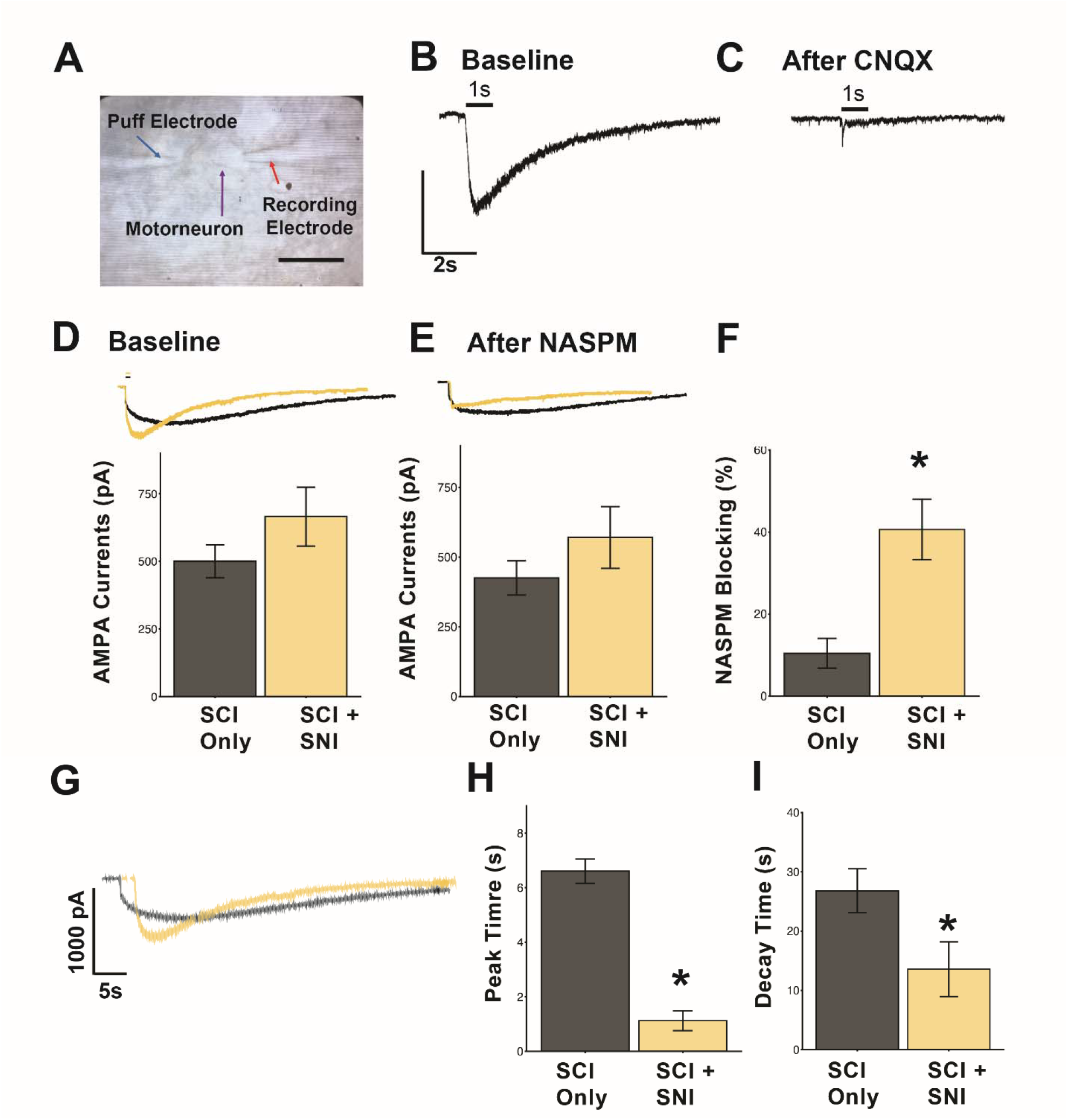
NASPM modulates the impact of nociceptive input (nerve injury) through the blockage of the GluA1 subunit of AMPA receptors. (A) Representative image of patch clamp procedure, showing puff and recording electrodes around motorneurons. NASPM, a selective CP-AMPAR antagonist showed a significant blocking effect on AMPA-induced currents in the motorneurons from both SCI and SCI+SNI groups. For the blocking percentage, the SCI+SNI group showed a larger effect than the SCI group. Representative traces and quantification of AMPA-induced currents (D) before and (E) after Naspm application (250 µM for 3 min) in motorneurons. Bars above the traces indicate the duration of AMPA application. (F) NASPM had the greatest blocking effect in the SCI Only group (7.2 ± 2%, n = 5 neurons from 5 mice) and significantly less blocking effect in neurons from mice with SCI + SNI (39.7 ± 6.8%, n = 7 neurons from 7 mice); *p = 0.01. (G) Representative traces of SCI Only and SCI + SNI to illustrate peak time and decay. Quantification of peak time (H) and peak delay (I) indicated significantly slower peak time and shorter decay time for the SCI + SNI group (p < 0.05). All data were expressed in Mean ± SEM.

AMPA puff-induced fast inward currents in all the recorded neurons from both SCI and SCI+SNI groups (**Figure 3D**). In the SCI group, the currents were 481.6 ± 66.2 pA (n = 12), while the SCI+SNI group demonstrated AMPA-induced currents at 607.9 ± 112.9 pA (n = 8), but no significant difference was discerned between groups (**Figure 3D**). Next, to identify how much AMPA currents were contributed by GluA2-containing AMPA currents, the specific CP-AMPAR antagonist NASPM was applied to the tissue bath to block the GluA1-containing AMPA currents. We found that after application of NASPM, there was no significant difference among SCI and SCI+SNI groups (425.6 ± 61.36 pA vs 425.3 ± 108.0 pA, respectively; n = 5-8 per group, p > 0.05, **Figure 3E**). Accordingly, we analyzed the blocking percentage of NASPM on AMPA currents, which demonstrated NASPM having an inhibitory effect in the SCI and SCI+SNI groups (7.2 ± 2.0% and 39.7 ± 6.8% reduction, respectively; n = 5-8 per group; **Figure 3F**).

### Distinct kinetics of AMPA-induced currents

Results from the NASPM application suggest that different forms of injury may alter the compositions of AMPA receptors. To test this, we compared the kinetics of AMPA-induced currents in SCI vs SCI+SNI. As shown in **Figure 3G**, there was a distinct difference in the currents, with the SCI group demonstrating a slower speed to reach the peak. In contrast, the SCI+SNI group shared similar demonstrated a faster speed. Therefore, we analyzed the time from the onset of the currents to the peak currents and found that, in the SCI group, the peak time is longer than the SCI+SNI group (6.6 ± 0.4 S vs. 1.6 ± 0.3 s, p < 0.001, n = 4-6; **Figure 3H**). Decay time showed similar changes, though less drastic (p < 0.05; **Figure 3I**). These findings confirmed our speculation that SCI paired with spared nerve injury produces a different composition of AMPA receptors on motorneurons than SCI alone.

### Broad range of aberrant peripheral input after SCI drives multidimensional AMPA receptor changes in ventral spinal cord synaptosomes

The above results indicate that a spared nerve injury model of polytraumatic SCI induces an AMPAR-mediated maladaptive plasticity. Given this, we considered whether this effect generalizes to a broader range of polytrauma models. Using data gathered from different types of aberrant stimulation below a complete SCI (e.g., electro-nociception, SNI, or hindlimb disuse/unloading), we carried out a large-scale analysis of all biomolecular protein endpoints. For each of these separate studies, near-IR quantitative Western blotting was performed on spinal cord synaptoneurosome isolates to assess four specific targets: GluA1, GluA2, pS831, and pS880, to assay AMPAR activity. Using a non-linear principal component analysis (NL-PCA), we uncovered a multidimensional pattern that accounts for 26.4% of the variance across GluA1, GluA2, pS831, and pS880 levels, as demonstrated in **Figure 4**. The resulting PC loading patterns revealed a pattern for PC1 in which all AMPAR protein targets loaded highly in the same direction, indicative of normal overall AMPAR plasticity. For PC2, GluA1 is increased, and GluA2 is decreased, indicating a shift toward CP-AMPAR expression (**Figure 4A**). The phosphorylation levels at PKC/CaMKII sites on AMPARs showed a similar pattern, with pS831 (GluA1) increasing and pS880 (GluA2) decreasing. This pattern indicates a CP-AMPAR maladaptive plasticity. Individual PC scores were then plotted on a biplot of PC1 (normal AMPAR plasticity) and PC2 (maladaptive plasticity) and colored by experimental condition to visualize the syndromic plasticity space (**Figure 4B**). Data were also filtered to reveal the unique position of SCI+Sham Nerve Injury and SCI+SNI groups in this proteomic space (**Figure 4C**). Planned comparison of PC2 (maladaptive plasticity) scores between the SCI and SCI+SNI groups showed a significant difference between groups, with SCI+SNI exhibiting higher scores on this maladaptive plasticity index (p < 0.05; **Figure 4D**). These results demonstrated that a broad range of nociceptive stimuli below the level of SCI induce maladaptive plasticity relative to their appropriate control groups through CP-AMPAR-mediated mechanisms.

**Figure 4.**
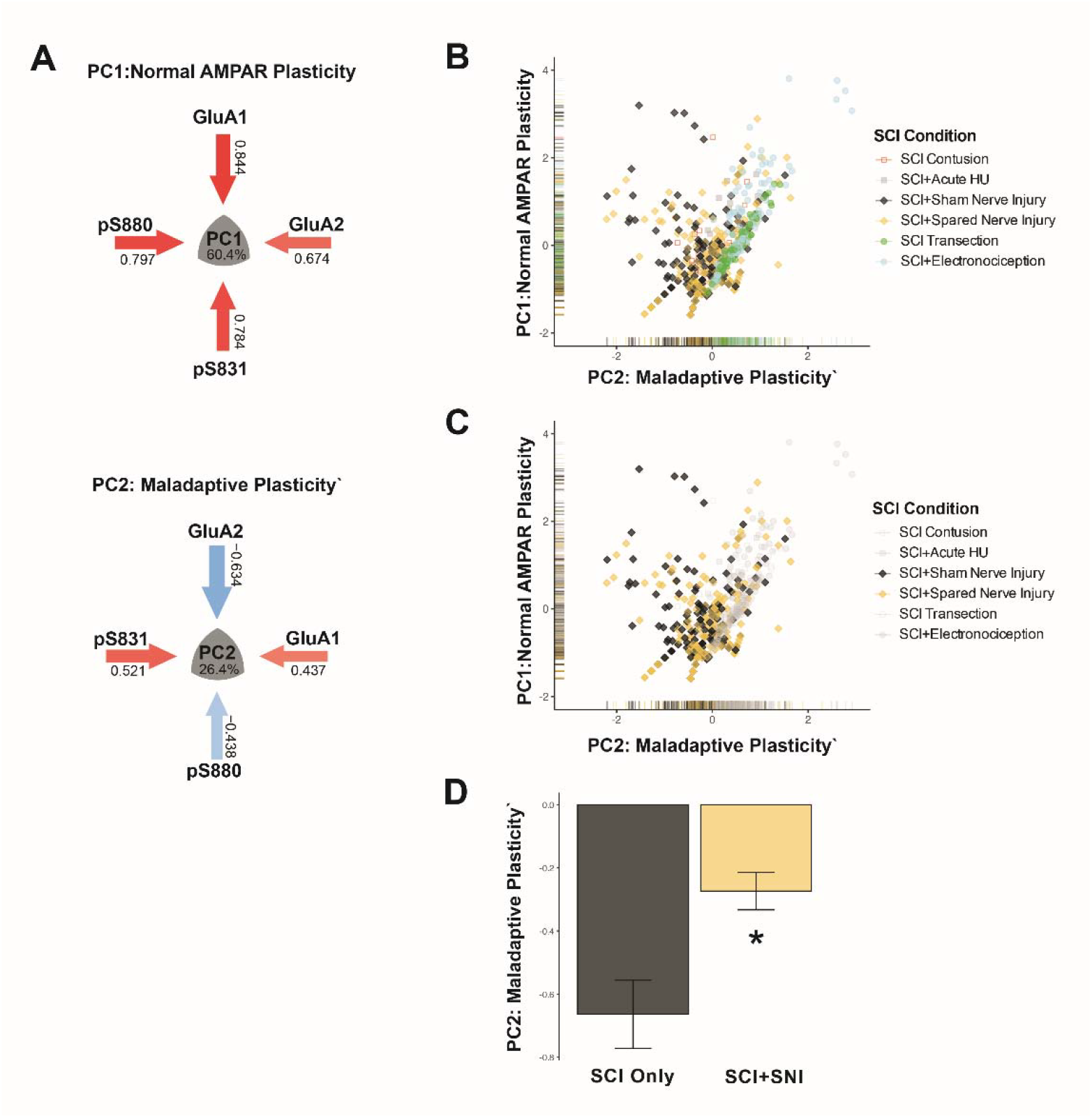
Large-scale analysis of all biomolecular protein endpoints pooled across different SCI models (contusion and complete spinal transection) and different forms of aberrant stimulation (electronociception, spared nerve injury, or hindlimb unloading conditions). (A) Non-linear principal component analysis (NL-PCA) uncovered a multidimensional pattern (principal component; PC) between GluA1, GluA2, pS831, and pS880 levels across acute electronociception/nerve injury preps (20 min post injury) and chronic hindlimb unloading (8 weeks post injury). Examination of the PC loading patterns (arrows) reveals PC1 (accounting for 60.4% of total variance), in which all AMPAR targets move in concert, indicative of normal AMPAR plasticity. PC2 revealed a pattern of increased GluA1 and reduced GluA2 (reflecting calcium-permeable AMPARs). Phosphorylation levels at PKC/CaMKII (‘calcium sensor’ kinase) sites on AMPARs mirrored this change, with pS831 (GluA1) increasing and pS880 (GluA2) decreasing. This pattern was identified as reflecting a maladaptive plasticity signature. B) Individual PC scores were plotted on a PC1/PC2 biplot and colored by experimental condition to visualize the syndromic plasticity space. C) Data were filtered to reveal the unique position of SCI+Sham Nerve Injury and SCI+SNI groups in this proteomic space. D) Hypothesis testing of SCI+SNI on the maladaptive plasticity component (PC2) revealed a significant increase in PC2 score relative to SCI+Sham Nerve Injury (* p < 0.05).

## Discussion

Our results demonstrate that a spared nerve injury model of polytraumatic SCI induces CP-AMPAR-mediated synaptic plasticity in spinal motor systems. These findings suggest that peripheral nerve injury acutely increases the insertion of CP-AMPARs to spinal synapses, which NASPM protected against in patch-clamp experiments, indicating the necessity of CP-AMPAR activity for maladaptive spinal plasticity. The results of this study corroborate previous findings, suggesting that peripheral injuries may be a factor in the inhibition of functional recovery following SCI.^7,16,54^ We and others have previously shown that this form of plasticity can be induced by a broad range of peripheral nociceptive/aberrant modalities and reflects a lasting maladaptive alteration that facilitates hypersensitivity/spasticity and impairs long-term locomotor recovery.^46,47,49,55^ Given that this mechanism of maladaptive plasticity is observed across a spectrum of peripheral inputs, overexpression of synaptic CP-AMPARs may reflect a common dysregulated compensatory response within the injured spinal cord. This finding will be an important avenue for further study, as these models of aberrant peripheral input, ranging from nerve injury to limb disuse, are seen widely in the clinical SCI population.^9–12^ Thus, CP-AMPARs may be a druggable therapeutic target for restoring beneficial synaptic plasticity after polytraumatic SCI.

## Methods

### Animals

Adult female C57BL/6 mice (Jackson Laboratories) were used for patch-clamp and western blot analyses. All procedures were conducted in accordance with the National Institute of Health Guide for the Care and Use of Laboratory Animals, and all efforts were made to minimize suffering and limit the number of animals used. All protocols were approved by the University Laboratory Animal Care Committee at the University of California, San Francisco.

### Surgeries

All animal subjects underwent a complete spinal cord transection immediately rostral to the ninth thoracic vertebra (T9). Animals were fully anesthetized with isoflurane gas (5%). Fur over the thoracic vertebra was shaved, and a 3 cm incision was made over T9. The tissue immediately rostral to T9 was cleared away with rongeurs, and the underlying spinal cord was exposed. An electrical heat cautery device was used to transect the spinal cord, and the resulting cavity was filled with Gelfoam (Harvard Apparatus). The incision was then closed using Michel clips (Fine Science Tools). Animals received a 2.5 ml intraperitoneal injection of 0.9% saline immediately following surgery and twice daily thereafter to ensure proper hydration. Bladders were expressed twice daily. Given the nociceptive nature of this study, no analgesics were given following complete spinal transection.

### Spared Nerve Injury

In this procedure, either the tibial or sural nerve is spared while the common peroneal and sural/tibial (depending on which was previously spared) nerve is tied off and severed. This model is summarized in **Figure 1**.

### Western Blot

Fresh-frozen (−80°C) spinal cords were subsequently rapidly thawed on a chilled petri dish at 4°C, and a 1 cm section of the lumbar enlargement was dissected. This section was then split horizontally. (Fig. 1A). The ventral hemicord (containing the ventral horn spinal gray matter) was then placed in a Dounce homogenizer filled with 200 m homogenization buffer (10 mM Tris, 30 mM sucrose, pH 7.5) containing phosphatase and protease inhibitors (Roche). Following tissue disruption, the whole homogenate was placed in an Eppendorf tube and spun at 5000 relative centrifugal force (rcf) for 5 min in a minicentrifuge at 4°C. This centrifugation procedure produced a supernatant (S1) and a nuclear pellet (P1). The S1 layer was removed and centrifuged a second time at 13,000 rcf for 30 min at 4°C, producing a modestly synaptoneurosomal-enriched pellet fraction (P2) that was subsequently used in Western blots (Fig. 1B).^56–58^ Total protein concentration was quantified using the Pierce BCA protein assay method. Each sample was diluted 1:2 in room-temperature Laemmli sample buffer, and 20 g of total protein per sample was loaded into separate lanes on a precast 10–20% Tris-HCl polyacrylamide gel (Bio-Rad). Samples were counterbalanced across the gel by treatment condition, and the experimenter was kept blind to the condition. A kaleidoscope ladder was loaded on each gel to confirm molecular weight. The gel was electrophoresed for 1 h at 100 V in SDS buffer (25 mM Tris, 192 mM glycine, 0.1% SDS, pH 8.3; BioRad). Protein was transferred to a nitrocellulose membrane in cold transfer buffer (25 mM Tris, 192 mM glycine, 20% methanol, pH 8.3). The membrane was blocked for 1 h in Odyssey Blocking Buffer (LiCor) containing 0.1% Tween 20, followed by an overnight incubation in primary antibody solution at 4°C. The primary antibody solution consisted of rabbit polyclonal antiphosphorylated serine 831 (1:200; Millipore) in Odyssey blocking buffer plus 0.05% Tween 20. Following incubation, the membrane was washed 4 times for 5 min with Tris-buffered saline containing 0.1% Tween 20 (TTBS), and incubated in a fluorescent-labeled secondary antibody (1:30K, Li-Cor IRDye 680 goat anti-rabbit in Odyssey blocking buffer plus 0.2% Tween 20) for 1 h in the dark. After 4 5-min washes in TTBS, followed by a 5-min wash in TBS. The membrane was immediately scanned on the Li-Cor Odyssey Infrared Imaging System using a 680 nm laser to detect protein bands. The membrane was then re-blocked and reincubated with mouse anti-GluA1 primary antibody (1:200, Millipore) in Odyssey blocking buffer plus 0.05% Tween 20. The membrane was then washed as before, incubated in a fluorescent-labeled secondary antibody (1:30K, Li-Cor IRDye 800 goat anti-mouse in Odyssey blocking buffer plus 0.2% Tween 20), washed as before, and rescanned with the 800 nm laser to reveal GluA1 protein bands. The same protocol was performed for serine 845 on GluA1, GluA2, and serine 880 on GluA2. Loading controls were performed as the last step on each multiplexed Western blot.

### Spinal Cord Slice Preparation and Whole Cell Patch Clamp Recording

The whole procedure was based on the previous publication with modest modification (Mitra, JNP, 2012). A total of 2 solutions were prepared for recording: dissection artificial cerebrospinal fluid (aCSF) and dissecting aCSF (DaCSF), which were used for slicing the spinal cord and recording aCSF (RaCSF) during the following patch clamp recording. For DaCSF, it was composed of (in mM) 191 sucrose, 0.75 K-gluconate, 1.25 KH2PO4, 26 choline bicarbonate (80% solution), 4 MgSO4, 1 CaCl2, 20 dextrose, 2 kynurenic acid sodium salt (Ascent Scientific, Bristol, UK), 1 (+)-sodium L-ascorbate, 5 ethyl pyruvate, and 3 myo-inositol. The DaCSF was freshly made and bubbled with carbogen on ice for 1 hr before usage.

After laminectomy, the spinal cord (L1-5 segments) was quickly dissected and transferred to a petri dish containing DaCSF bubbled with carbogen on ice. The meninges and nerve roots were removed using scissors under a dissection microscope. For spinal cords from the SCI + SNI group, the contralateral side (right) of the spinal cords was cut on the edge to be differentiated. Then the spinal cord was placed on a clean petri dish and immediately covered with warm, low-melting agarose (1.5% in DaCSF, 37 °C), followed by cooling on ice until the agarose hardened. Then the agarose gel block was trimmed and sectioned on a vibratome (VTS 1000, Leica, Germany) with a thickness of 350 µm. The whole slicing procedure was conducted in ice cold DaCSF bubbled with carbogen. After slicing, the slices were collected into a slice holder and incubated in 35 °C DaCSF for 30 min, followed by incubation in RaCSF at room temperature for the remainder of the experiment.

Neurons were visualized with a 40X water-immersion objective using a microscope (FN-600, Nikon, Japan) equipped with infrared differential interference contrast optics. The image was detected with an infrared-sensitive CCD (IR-1000, Dage MTI, USA) and displayed on a monochrome video monitor. Motorneurons were easily identified by their size (> 40 µm), shape (pyramidal), and position (Ventral horn). Patch pipettes were pulled from borosilicate glass capillaries (BF150-86-10, Sutter Instruments) on a puller (P97, Sutter Instruments, Novato, CA, USA).

Two types of internal solutions were used during patch-clamp recordings. For action potential and α-amino-3-hydroxy-5-methyl-4-isoxazolepropionic acid (AMPA) induced inward currents recording, the following internal solution was used (in mM): 140 K-Gluconate, 2 MgCl_2_, 10 HEPES, 2 Mg-ATP, 0.5 Na_2_GTP, pH□ = □7.4. Osmolality was adjusted to 290–300 mOsm. For recording of miniature excitatory postsynaptic currents (mEPSC) or miniature inhibitory postsynaptic currents (mIPSC), Cesium containing internal solution was used, which contained (in mM): Cs_2_SO4, 110; CaCl_2_, 0.5; MgCl_2_, 2; EGTA, 5; HEPES, 5; tetraethylammonium (TEA), 5; with pH adjusted to 7.2–7.4 by CsOH, and had an osmolarity of 290–300 mOsm.

After a giga seal was established, the membrane was broken, and neurons with a resting membrane potential (RMP) below −50 mV were selected for further study. The access resistance was continuously monitored at 10–20 MΩ, and data were discarded if it changed by more than 15% during an experiment.

For all agonists induced currents, the neurons were clamped at −70 mV. The methods for measuring rheobase and membrane threshold were described in detail in our previous publication.^59^ Briefly, a series of currents starting at −0.1 nA with increments of 0.05 nA was administered until the first action potential was generated to measure the membrane threshold. A 500 ms depolarizing ramp (2000 pA/s) was administered to measure the membrane threshold.

For mEPSC and mIPSC recordings, the neurons were clamped at −70 mV and 0 mV, respectively. Tetrodotoxin (TTX, 10 nM) and bicuculline (10 µM) were bath applied for mEPSC recording; while TTX, CNQX (10 µM), and APV (50 µM) were bath applied for mIPSC recording.

All recordings were acquired with an Axon 200B amplifier (Molecular Devices, Sunnyvale, CA, USA). Data were acquired with a Digidata 1440 A acquisition system (Molecular Devices) and pCLAMP 10.2 software (Molecular Devices). Signals were low-pass filtered at 5 kHz, sampled at 10 kHz, and analyzed offline.

### Drug application

All drugs used in the □ex vivo □experiments were purchased from Tocris (Tocris, Minneapolis, MN, USA). Drugs were dissolved in ultra-pure deionized water as stock solutions and refrigerated for later use. The stock solution was diluted to the desired concentration with RaCSF immediately before use. AMPA (100 µM) was applied focally by pressure ejection via a micropipette connected to a Picrospitzer II (puff at 1–2 psi, General Valve, USA) using nitrogen as a gas. The pipette tip was located approximately 50 µm from the recorded neuron, ensuring that drugs reached all parts of the neuron.

CNQX (10 µM), APV (50 µM), TTX (1 nM), and NASPM (50 µM) were bath-applied for at least 5 minutes to test their blocking effects. In order to test the blocking effect of receptor antagonists, the responses induced by each agonist before antagonist application were set as 100%, and the currents after antagonist application were expressed as the percentage of the previous response. The tau of decay is calculated as the time from 100% of the peak value to 50% of the peak value.

### Statistical analyses

Western blot data were analyzed using a linear mixed model, with technical replicates across western blot gels treated as a random factor within each subject, and the ratio of target protein to loading control used as the outcome measure.^53^ For patch clamp experiments, a paired Student’s □ t-test was used to compare agonist induced inward currents and antagonists induced inward currents. Principal components analysis was run using the *syndRomics* package.^60^ All bar graph results are presented as the mean ± SEM.

